# *Trans*-acting genetic variation affects the expression of adjacent genes

**DOI:** 10.1101/2020.10.05.327130

**Authors:** Krisna Van Dyke, Gemechu Mekonnen, Chad L. Myers, Frank W. Albert

**Affiliations:** Department of Genetics, Cell Biology, and Development, University of Minnesota, Minneapolis, MN, U.S.A

## Abstract

Gene expression differences among individuals are shaped by *trans*-acting expression quantitative trait loci (eQTLs). Most *trans*-eQTLs map to hotspot locations that influence many genes. The molecular mechanisms perturbed by hotspots are often assumed to involve “vertical” cascades of effects in pathways that can ultimately affect the expression of thousands of genes. Here, we report that *trans*-eQTLs can affect the expression of adjacent genes via “horizontal” mechanisms that extend along a chromosome. Genes affected by *trans*-eQTL hotspots in the yeast *Saccharomyces cerevisiae* were more likely to be located next to each other than expected by chance. These paired hotspot effects tended to occur at adjacent genes that show coexpression in response to genetic and environmental perturbations. Physical proximity and shared chromatin state, in addition to regulation of adjacent genes by similar transcription factors, were independently associated with paired hotspot effects. The effects of *trans*-eQTLs can spread among neighboring genes even when these genes do not share a common function. This phenomenon could result in unexpected connections between regulatory genetic variation and phenotypes.

## Introduction

Genetic variation among individuals influences phenotypic traits (Lynch & Walsh, 1998). Many of the DNA variants that shape phenotypes do so by altering gene expression (Albert & Kruglyak, 2015; Gusev et al., 2014; Maurano et al., 2012; Signor & Nuzhdin, 2018; Zheng et al., 2011). Genomic regions that contain variants that influence the abundance of the RNA of a given gene are called expression quantitative trait loci (eQTLs) (Albert & Kruglyak, 2015; Brem, 2002; W. Sun & Hu, 2013). eQTLs can be classified according to their location in the genome relative to the genes they affect, as well as by their mechanism of action. Local eQTLs alter the expression of genes in their physical vicinity, usually by perturbing *cis*-acting regulatory mechanisms. By contrast, distant eQTLs act in *trans* by changing the abundance, cellular localization, or activity of a diffusible factor. These factors can act on genes throughout the genome, and most *trans*-eQTLs influence genes that are distant from the eQTL, typically on different chromosomes.

*Trans*-acting variation has been extensively studied in crosses in model organisms (Brem, 2002; Cesar et al., 2018; Everett et al., 2020; Hubner et al., 2005; Lewis et al., 2014; Orozco et al., 2012; Rockman et al., 2010; Smith & Kruglyak, 2008; West et al., 2007). For example, genetic mapping of mRNA levels in 1,012 recombinant offspring from a cross between a laboratory strain (BY, a close relative of the genome reference strain S288C) and a vineyard isolate (RM) of the yeast *Saccharomyces cerevisiae* yielded tens of thousands of eQTLs. These eQTLs accounted for the majority of genetic variation in mRNA abundance (a median across genes of 72%), suggesting that the dataset comprises the great majority of eQTLs with at least modest effect sizes. Almost all genes were influenced by multiple *trans*-eQTLs. The summed *trans*-eQTL effects accounted for 2.6-fold more expression variation than those of local eQTLs (Albert et al., 2018). In agreement with similar estimates in human populations (Grundberg et al., 2012; Wright et al., 2014), the BY / RM cross shows that *trans*-eQTLs are the major source of genetic variation in gene expression.

*Trans*-eQTLs tend to cluster at certain locations in the genome. These *trans*-eQTL “hotspots” can affect the expression of large numbers of genes, and have been observed in yeasts (Brem, 2002; Clément-Ziza et al., 2014; Smith & Kruglyak, 2008), nematodes (Rockman et al., 2010), plants (Fu et al., 2009; West et al., 2007), rodents (Bystrykh et al., 2005; Chesler et al., 2005; Langley et al., 2013; Orozco et al., 2012), and cattle (Cesar et al., 2018), and may also exist in humans (Brynedal et al., 2017). In the BY / RM yeast cross, over 90% of the *trans*-eQTLs mapped to 102 hotspot regions (Albert et al., 2018). Several of these hotspots have been resolved to single causal genes or variants (Brem, 2002; Lutz et al., 2019; Smith & Kruglyak, 2008; Yvert et al., 2003; Zhu et al., 2008). The BY / RM hotspots can affect hundreds and in some cases thousands of genes.

Identifying the mechanistic connection between an eQTL hotspot and the genes it affects in *trans* can be challenging. Typically, the causal variant or variants in the hotspot are assumed to alter the sequence or abundance of a protein product, such as a transcription factor, that then acts on functionally related genes. In turn, the changes in gene expression of these genes are propagated through cellular networks to indirectly affect many other genes. In this paper, we refer to this type of mechanistic connection as “vertical propagation”.

The expression of eukaryotic genes is partially shaped by their location on the chromosome. Specifically, adjacent genes tend to show correlated expression changes in response to a broad range of stimuli (Cohen et al., 2000; Hurst et al., 2004; Kustatscher et al., 2017; Michalak, 2008; M. Sun & Zhang, 2019). In part, this coexpression is due to clustering of genes with similar function and regulation by transcription factors (TFs) (Allocco et al., 2004; Arnone et al., 2012; Cohen et al., 2000). In addition, shared local chromatin states that encompass adjacent genes may facilitate coexpression even for genes that do not share a common function and that are not regulated by the same TFs (Arnone et al., 2014; Batada et al., 2007; Ebisuya et al., 2008; M. Sun & Zhang, 2019). Coexpression of neighboring genes decreases with physical distance (Cohen et al., 2000; Quintero-Cadena & Sternberg, 2016). In addition to chromatin state, non-specific promiscuous effects of *cis*-regulatory elements have been proposed to contribute to this distance-dependent coexpression (Cohen et al., 2000; Quintero-Cadena & Sternberg, 2016).

Here, we asked if the effects of *trans*-eQTL hotspots propagate “horizontally” along a chromosome. We made use of the extensive set of *trans*-eQTLs identified in the BY / RM cross (Albert et al., 2018) to show that *trans*-eQTL hotspots affect the expression of adjacent genes more frequently than expected by chance. This effect occurs primarily at adjacent genes that show correlated gene expression responses to a broad variety of perturbations. Paired hotspot effects can be partially explained by regulation of adjacent genes by similar transcription factors. However, paired hotspot effects were independently associated with the physical proximity of the genes in a pair as well as similar chromatin states, suggesting that the effects of *trans*-eQTLs can extend from an affected gene to adjacent genes via “horizontal” mechanisms that spread along the chromosome. This work demonstrates the existence of an additional mode by which *trans*-acting regulatory variation can affect gene regulatory networks and cell biology.

## Results

### *Trans*-eQTL hotspots affect more adjacent gene pairs than expected by chance

We analyzed eQTLs mapped in the BY and RM cross (Albert et al., 2018). Briefly, the dataset comprises 36,498 eQTLs, of which 33,529 act in *trans*. These *trans*-eQTLs cluster at 102 hotspot locations, which affect 12 to 4,093 of 4,912 verified open reading frames (ORFs) with detectable expression. Genes are affected by a median of eleven hotspots, and almost all genes (98%) are affected by at least one hotspot.

We asked whether hotspots affected genes located next to each other on the chromosome more often than expected by chance. To address this question, we focused on “doublets”, which we defined as pairs of adjacent verified ORFs whose mRNA abundances were both increased or both decreased by a given hotspot. Genes close to chromosome ends were excluded from the analysis (Methods). For each hotspot, we constructed 10,000 permutations by randomly assigning the observed effects of the hotspot to genes, irrespective of their location in the genome. This permutation strategy preserved the number and magnitude of hotspot effects, but broke any relationship between hotspot effects and a gene’s location relative to its neighbors. We compared the number of observed doublets to the distribution of doublets in the hotspot-specific permutations.

Out of the 102 hotspots, 91 displayed more doublets than their matched permutation median. This number was significantly higher than expected by chance (Binomial Test, p < 2.2×10^−16^). At 58 hotspots, the observed number of doublets exceeded that seen in 95% of the permutations (Fig 1, Suppl Table 1, Suppl Fig 1), which was more than the five hotspots expected to reach this threshold by chance (Binomial Test, p < 2.2×10^−16^). After Bonferroni correction, 17 hotspots exhibited a significant excess of doublets (Table 1, Fig 1, Suppl Table 1, Suppl Fig 1). Coexpression of adjacent genes might occur due to shared promoter sequences when the genes are transcribed in divergent orientations. For example, in the yeast genome, genes in the ribosome and rRNA biosynthesis regulons tend to occur in adjacent, divergently expressed pairs that show highly correlated expression (Arnone et al., 2014; Arnone & McAlear, 2011; Eldabagh et al., 2018; Kraakman et al., 1989; Wade et al., 2006). To test if the excess of paired hotspot effects was only due to gene pairs with divergent orientation, we excluded all divergent gene pairs from our permutation analysis. The number of doublets remained significantly higher than expected by chance (Table 1, Suppl Table 1).

**Figure 1:**
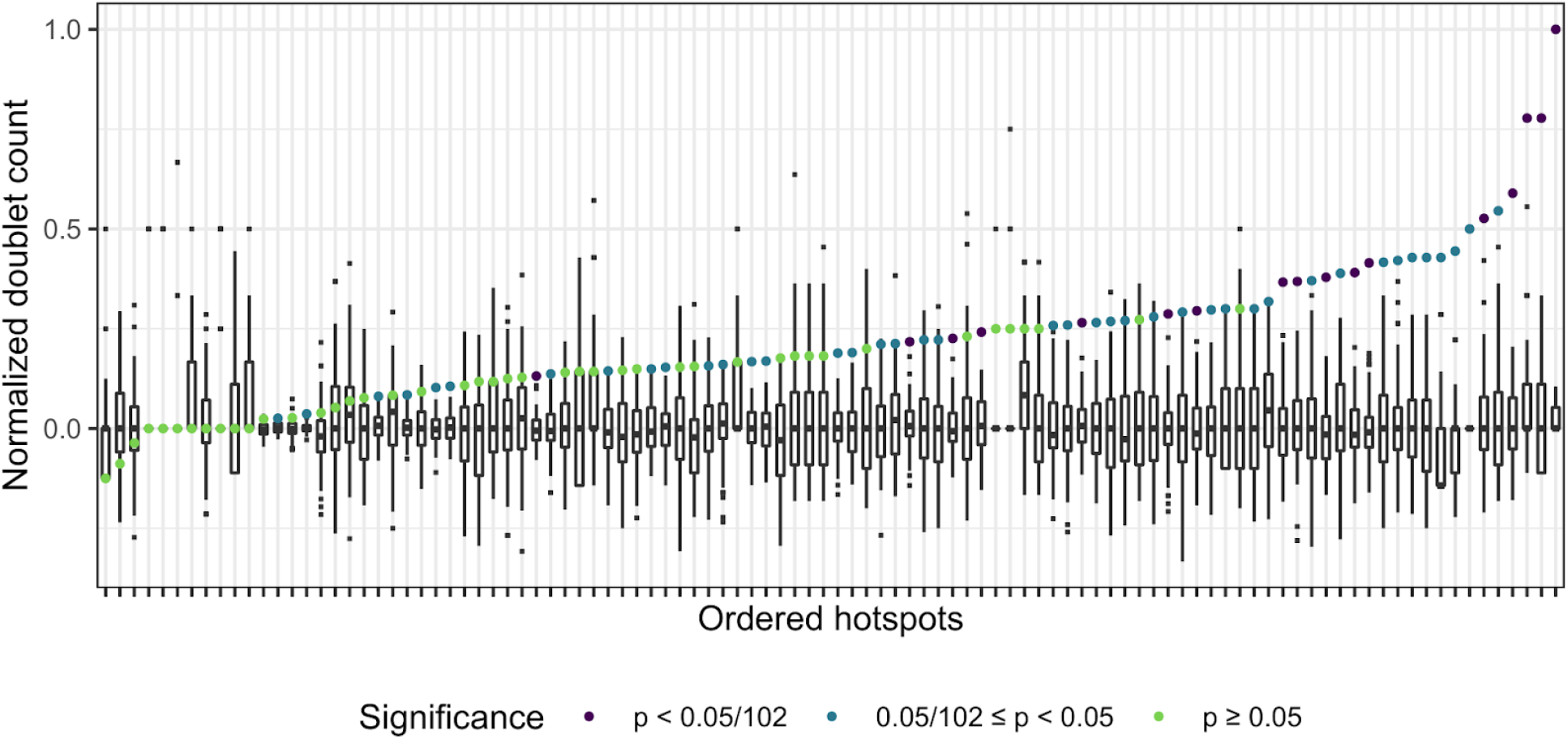
The number of doublets in real data (colored circles) are plotted against the number of doublets in permuted data (white boxes). Boxes extend from the 25th to 75th percentile. Whiskers extend from each box to largest or smallest values within 1.5 times the difference in the 25th to 75th percentile. Data beyond the whiskers are considered outliers and plotted as small black squares. Circles are colored by significance of the excess of doublets at the given hotspot. To aid visualization, the doublet count for each hotspot and its matched permutations were normalized by subtracting the median and dividing by the maximum value. See Suppl Fig 1 for non-normalized results. Hotspots are ordered along the x-axis based on the value of the normalized doublet count in the real data.

**Table 1:** Enrichment of adjacent genes affected by *trans*-eQTL hotspots

| | Number of hotspots that pass multiple test correction* | p-value** for hotspots that pass multiple test correction | Number of hotspots with $p < 0.05$ | p-value** for hotspots with $p < 0.05$ | Number of hotspots that exceed median permutation n | p-value** for hotspots that exceed median permutation n |
| --- | --- | --- | --- | --- | --- | --- |
| **Doublets** | 17 | $5 \times 10^{-38}$ | 58 | $6 \times 10^{-48}$ | 91 | $4 \times 10^{-17}$ |
| **Doublets without divergent pairs** | 17 | $5 \times 10^{-38}$ | 57 | $1 \times 10^{-46}$ | 90 | $3 \times 10^{-16}$ |
| **Triplets** | 7 | $1 \times 10^{-13}$ | 30 | $2 \times 10^{-15}$ | 66 | 0.002 |
| **Quadruplets** | 4 | $2 \times 10^{-7}$ | 18 | $3 \times 10^{-6}$ | 40 | 0.99 |
| **Quintuplets** | 2 | 0.002 | 12 | 0.005 | 19 | 1 |
| **Sextuplets** | 1 | 0.049 | 3 | 0.89 | 10 | 1 |

\* The multiple-test corrected p-value was obtained by dividing $p = 0.05$ by the number of hotspots (102).
\*\* P-values were computed using a binomial test of the alternative hypothesis of a greater number of hotspots than expected by chance. The number of hotspots expected by chance are 0.05 hotspots with multiple testing correction (i.e., $0.05 / 102$ ), 5.1 hotspots at $p < 0.05$ , and 51 hotspots at $p < 0.5$ .

The number of sets of three adjacent genes affected by hotspots in the same direction was also higher than expected by chance (“triplets” in Table 1; Suppl Table 1), as were sets of four adjacent genes (“quadruplets”) and sets of five adjacent genes (“quintuplets”), albeit at weaker significance. No enrichment was found for sets of six adjacent genes (“sextuplets” in Table 1). Thus, the mechanisms causing paired hotspot effects extend beyond immediate gene neighbors and eventually taper off as a function of distance. Overall, these results show that the genes affected by hotspots are not randomly located throughout the genome, suggesting that the effects of *trans*-acting variation may be shaped by spatial patterns of gene regulation along chromosomes.

### Gene coexpression patterns are shared across diverse transcriptome datasets

*Trans*-eQTLs could affect the expression of adjacent genes via mechanisms that lead to gene coexpression (Brown et al., 2013; Michalak, 2008; M. Sun & Zhang, 2019). To quantitatively test this hypothesis, we first examined coexpression patterns in eleven transcriptome datasets (Fig 2) (Albert et al., 2018; Brem & Kruglyak, 2005; Fleming et al., 2002; Hughes et al., 2000; Klevecz et al., 2004; Knijnenburg et al., 2009; Lenstra et al., 2011; Myers et al., 2019; Sameith et al., 2015; Schurch et al., 2016; Simola et al., 2010). Each dataset comprised multiple genome-wide transcriptome experiments (n = 32 – 1,012, median = 162) collected in yeast strains exposed to a wide range of perturbations, including engineered gene deletions and natural genetic variation, environmental stressors such as chemical treatments, and deprivation of specific nutrients. These datasets were independently collected by different laboratories using various microarray or RNA-seq techniques. We also included the expression data from which the hotspots studied here were identified (Albert et al., 2018). To avoid possible circularity in these data, in which the perturbations caused by the hotspots could induce correlations among adjacent genes, we used linear models to remove the effects of eQTLs on segregant expression data (Methods).

**Figure 2:**
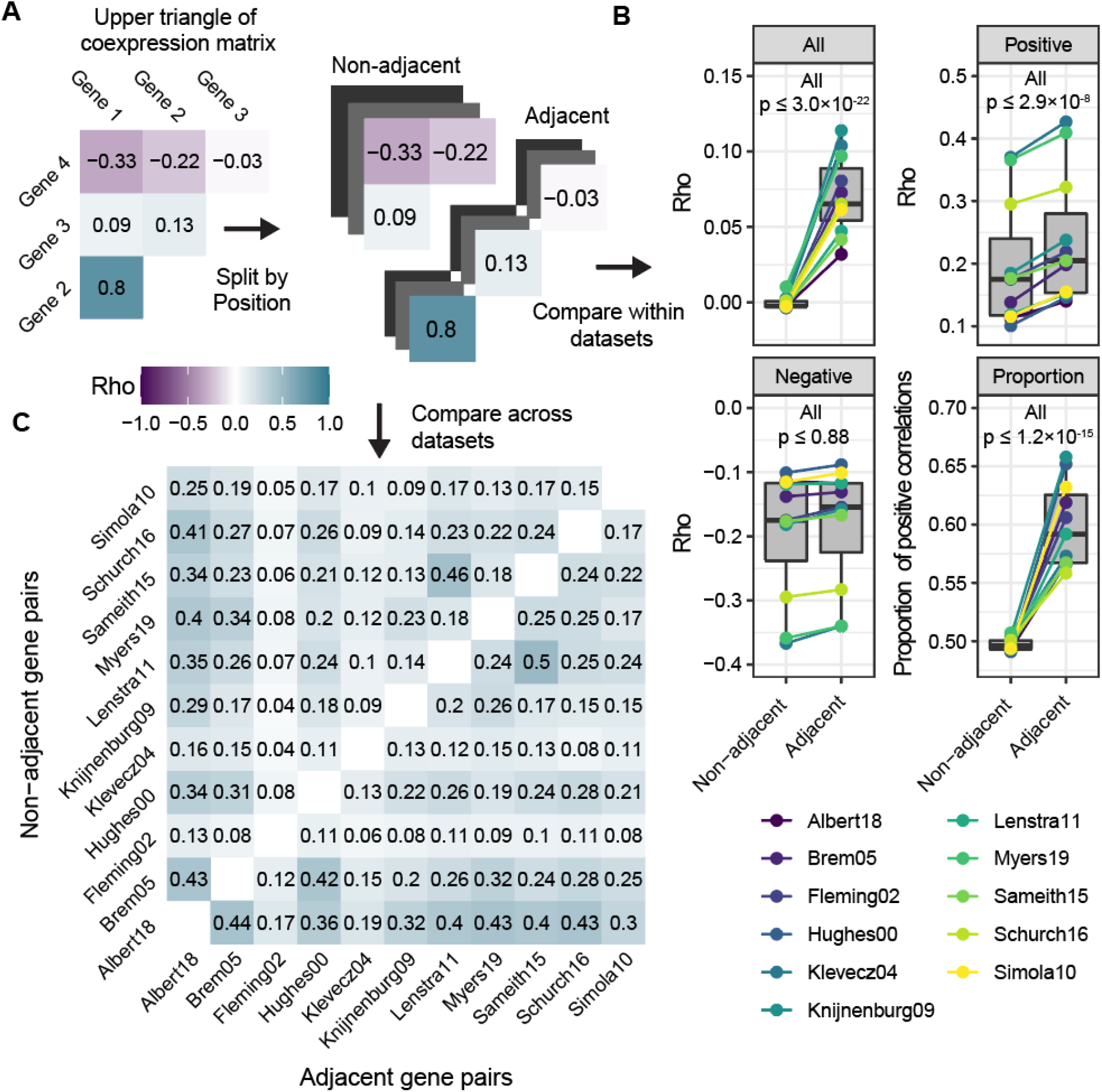
Coexpression analysis. (A) Schematic illustrating the separation of gene-gene coexpression matrices by adjacent versus non-adjacent genes. (B) Comparison of coexpression for adjacent genes versus non-adjacent genes in each of the eleven datasets, which are indicated by different colors. The four panels are, clockwise from top left: coexpression values (Rho) for all gene pairs, gene pairs with positive coexpression, gene pairs with negative coexpression, and the proportion of gene pairs with positive coexpression. (C) Correlations between coexpression datasets for neighboring vs non-neighboring genes. All correlations between datasets were statistically significant (p ≤ 8.6 × 10^−5^). The color scale is the same as in panel (A), and rho values are indicated in each cell.

To measure gene-gene coexpression in each of these transcriptome datasets, we built Spearman rank-based correlation matrices (Fig 2A). In all datasets, gene pairs showed a broad range of coexpression values, ranging from strongly positive to strongly negative correlations (Suppl Fig 2). Adjacent genes were more strongly correlated than non-adjacent genes in each dataset (p ≤ 3×10^−22^, Wilcoxon test, Fig 2B), consistent with earlier reports (Batada et al., 2007; Cohen et al., 2000; Ebisuya et al., 2008; Michalak, 2008; Raj et al., 2006). This overall correlation among adjacent genes was mostly driven by stronger positive as opposed to negative correlations, and there was a significant excess of positive correlations among adjacent compared to non-adjacent genes (Fig 2B).

Beyond these general patterns, we asked if the coexpression values for individual pairs of genes are quantitatively similar between the eleven different datasets. To do so, we computed the correlation between the coexpression relationships observed in each pair of datasets for either adjacent genes (Fig 2C, lower triangle) or non-adjacent genes (Fig 2C upper triangle). Both exhibited significant non-zero correlation in all comparisons (adjacent: p < 9×10^−5^; non-adjacent: p < 2.2×10^−16^). The correlation of coexpression of adjacent gene pairs between datasets was stronger than that of non-adjacent gene pairs (Median Rho = 0.24 vs 0.18, paired Wilcoxon test p < 2.2×10-16). These results show that coexpression relationships are broadly consistent across diverse environmental contexts and perturbations, especially for adjacent genes.

### *Trans*-eQTL hotspots tend to affect adjacent gene pairs with correlated expression

We tested if paired effects of *trans*-eQTL hotspots tend to occur at coexpressed gene pairs. To do so, we summarized hotspot effects on gene pairs across all hotspots into a score (Methods). Positive scores indicated that both genes in a pair are affected in the same direction (either both genes have increased or decreased expression) by multiple hotspots. Negative scores indicated pairs in which both genes were affected in opposite directions by multiple hotspots. A score near zero indicated pairs that were targeted by multiple hotspots but without a preference for the same or opposite direction, or gene pairs that were not affected by many hotspots. Adjacent gene pairs had higher scores (Wilcoxon test, p ≤ 1.1×10^−16^) and a higher proportion of positive scores (Test of proportions, p ≤ 8×10^−7^) than non-adjacent gene pairs, showing that this metric captured pairing of hotspot effects.

We compared these summarized hotspot effects to each of the eleven coexpression matrices, separately for adjacent genes and non-adjacent genes. Paired hotspot effects were significantly correlated with coexpression in all eleven datasets (Fig 3A). This agreement was present for adjacent genes (median of eleven Rhos = 0.21, all p < 2.2×10^−16^) and for non-adjacent genes (median of eleven Rhos = 0.18, all p < 2.2×10^−16^, Fig 3A), with stronger agreement for adjacent genes (paired Wilcoxon test: p = 5 ×10^−4^). The correlation between paired hotspot effects and coexpression arose primarily, but not exclusively, from gene pairs with stronger coexpression (Fig 3B). Thus, hotspots tend to affect both genes in pairs of genes that show stronger coexpression, particularly among adjacent genes. A parsimonious explanation for this agreement is that the molecular processes that cause coexpression are also responsible for spatial coupling of the effects of *trans*-eQTL hotspots.

**Figure 3.**
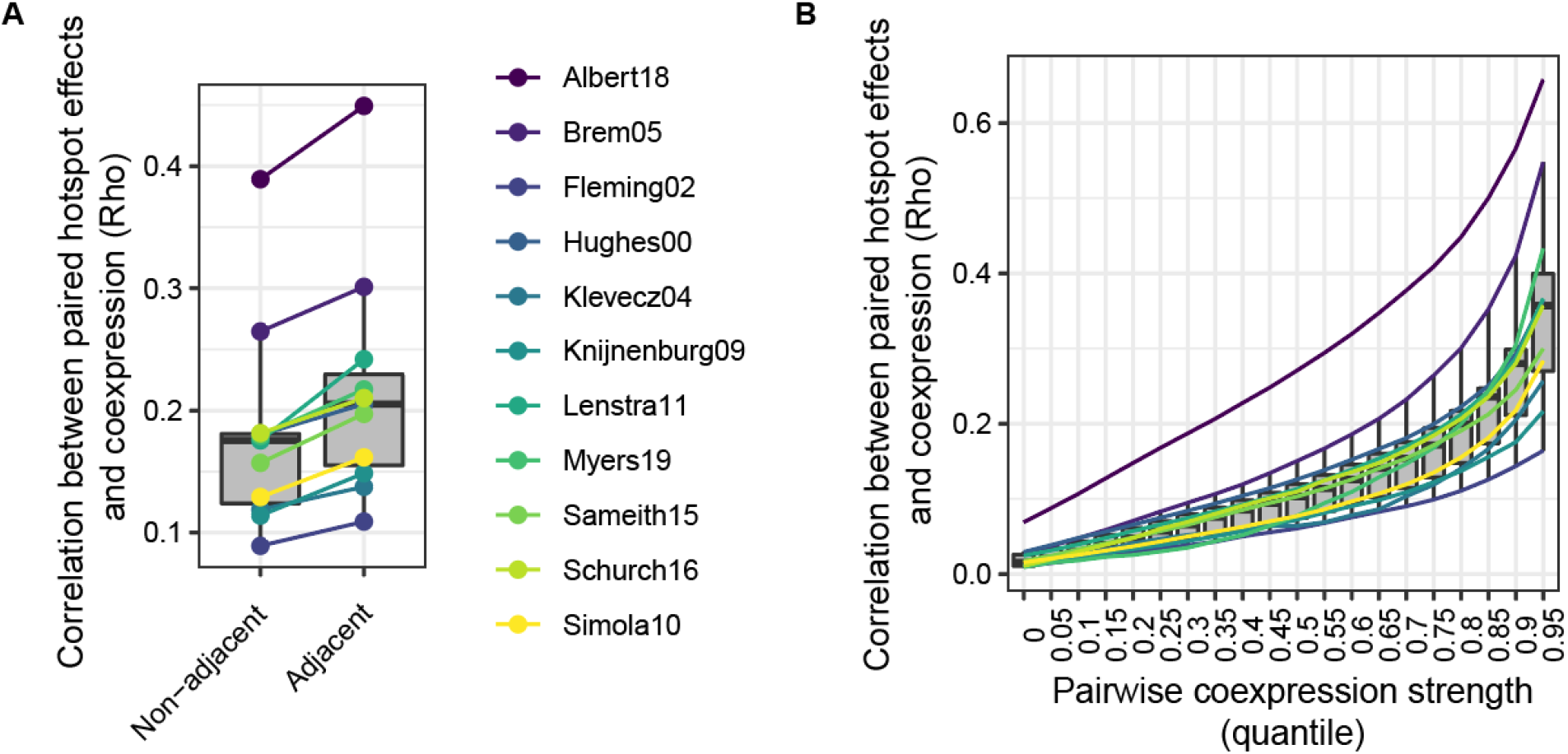
Agreement between paired hotspot effects and gene-gene coexpression. (A) Agreement between paired hotspot effects and the eleven coexpression datasets (shown in different colors), for non-adjacent and adjacent gene pairs. (B) Agreement between paired hotspot effects and coexpression datasets as a function of increasing coexpression. All Rho values, in all bins and for all datasets, were significantly different from zero (p ≤ 1.3×10^−16^).

### Paired hotspot effects on adjacent genes are driven by shared regulation by transcription factors, physical proximity, and similar chromatin dynamics

We asked which mechanisms might drive paired hotspot effects on adjacent genes. Specifically, we explored molecular features previously identified to affect adjacent gene coexpression, including regulation by similar sets of transcription factors, shared chromatin state, orientation of the genes relative to one another, and the distance between the genes. To relate hotspot effects on adjacent genes to these features, we counted the number of hotspots that affected both genes in a pair in the same direction. These counts ranged from zero to 41 (median = 2), with higher values for adjacent compared to non-adjacent genes (p = 1.1×10^−14^).

In *S. cerevisiae*, adjacent genes tend to be regulated by similar sets of transcription factors (Hershberg et al., 2005), likely because genes with similar functions tend to be located together in the genome (see below and (Cohen et al., 2000; Eldabagh et al., 2018)). To examine whether paired hotspot effects may result from shared regulation by transcription factors, we gathered annotated transcription factor binding sites (TFBS; (Monteiro et al., 2020)) in the intergenic region upstream of each gene and calculated the similarity of these TFBS annotations among all gene pairs. Adjacent genes had higher similarity in their TFBS profiles than non-adjacent genes (Median 0.294 vs 0.275, Wilcoxon test p < 2.2×10^−16^), as expected (Hershberg et al., 2005). TFBS similarity was weakly but significantly correlated with hotspot effect pairing for both adjacent genes (Rho = 0.033, p = 0.02) and non-adjacent genes (Rho = 0.027, p < 2.2×10^−16^). Thus, some paired hotspot effects are due to regulation of adjacent genes by shared transcription factors. This mechanism is consistent with traditional vertical propagation of hotspot effects to transcriptional target genes. However, the weak magnitude of the correlation between TFBS similarity and paired hotspot effects suggests that other mechanisms play a role.

A second possible mechanism that could create paired hotspot effects is shared chromatin state across neighboring genes, such that alterations to chromatin at one gene spread outward and also affect the other gene (Arnone et al., 2014; Brown et al., 2013; Raj et al., 2006). We first considered two independent measures of nucleosome occupancy across the bodies of adjacent genes (Chereji et al., 2018; Schep et al., 2015), but neither was correlated with paired hotspot effects (Chereji mean occupancy across adjacent gene bodies: p = 0.70; Chereji similarity in occupancy of adjacent genes: p = 0.06, Schep mean occupancy, p = 0.16; Schep similarity in occupancy, p = 0.36). These chromatin data, which were measured in normally growing, isogenic cultures, may not well reflect chromatin changes in response to a perturbation, such as a *trans*-eQTL. Therefore, we obtained data on 26 chromatin marks measured as a time course during the response to exposure to diamide, a stressful stimulus (Weiner et al., 2015). We focused on “+1” nucleosomes because of their important roles in transcriptional regulation (Jansen & Verstrepen, 2011; Rando & Winston, 2012). We computed two metrics: 1) similarity in chromatin state at baseline, which we defined as the correlation between the levels of chromatin marks on the +1 nucleosomes of adjacent genes before application of diamide, and 2) similarity in dynamic chromatin change, which we defined as the correlation between the *change* in +1 nucleosome chromatin marks of adjacent genes following cellular stress. Baseline chromatin state similarity showed no relationship with paired hotspot effects (p = 0.17). By contrast, similarity in chromatin dynamics showed a weak but significant association (rho = 0.03, p = 0.02).

Third, we compared genes expressed in different orientations (divergent, tandem, and convergent). Analysis of variance (ANOVA) showed a significant association between orientation and the number of paired hotspot effects (p = 2×10^−10^). Divergent gene pairs showed the most hotspot effect pairing, followed by pairs oriented in tandem and convergent pairs (Suppl Fig 3; Wilcoxon tests: divergent vs. tandem p = 0.014; divergent vs. convergent p = 9×10^−8^; tandem vs. convergent = 2×10^−4^).

Fourth, we asked how physical distance related to paired hotspot effects. Distance may be a proxy for a number of molecular mechanisms, including promiscuous effects of *cis*-regulatory elements that decay as a function of distance (Quintero-Cadena & Sternberg, 2016). While adjacent genes are all close to each other by definition, the distance between the start codons of the verified ORFs we analyzed here ranges from 46 bp to 19,141 bp, with a median of 1,924 bp (Fig 4A). We found that paired hotspot effects tended to occur at adjacent genes that were closer to each other (Rho = 0.05, p < 2.6×10^−6^; closeness was quantified as the log of the inverse of distance in bp).

**Figure 4:**
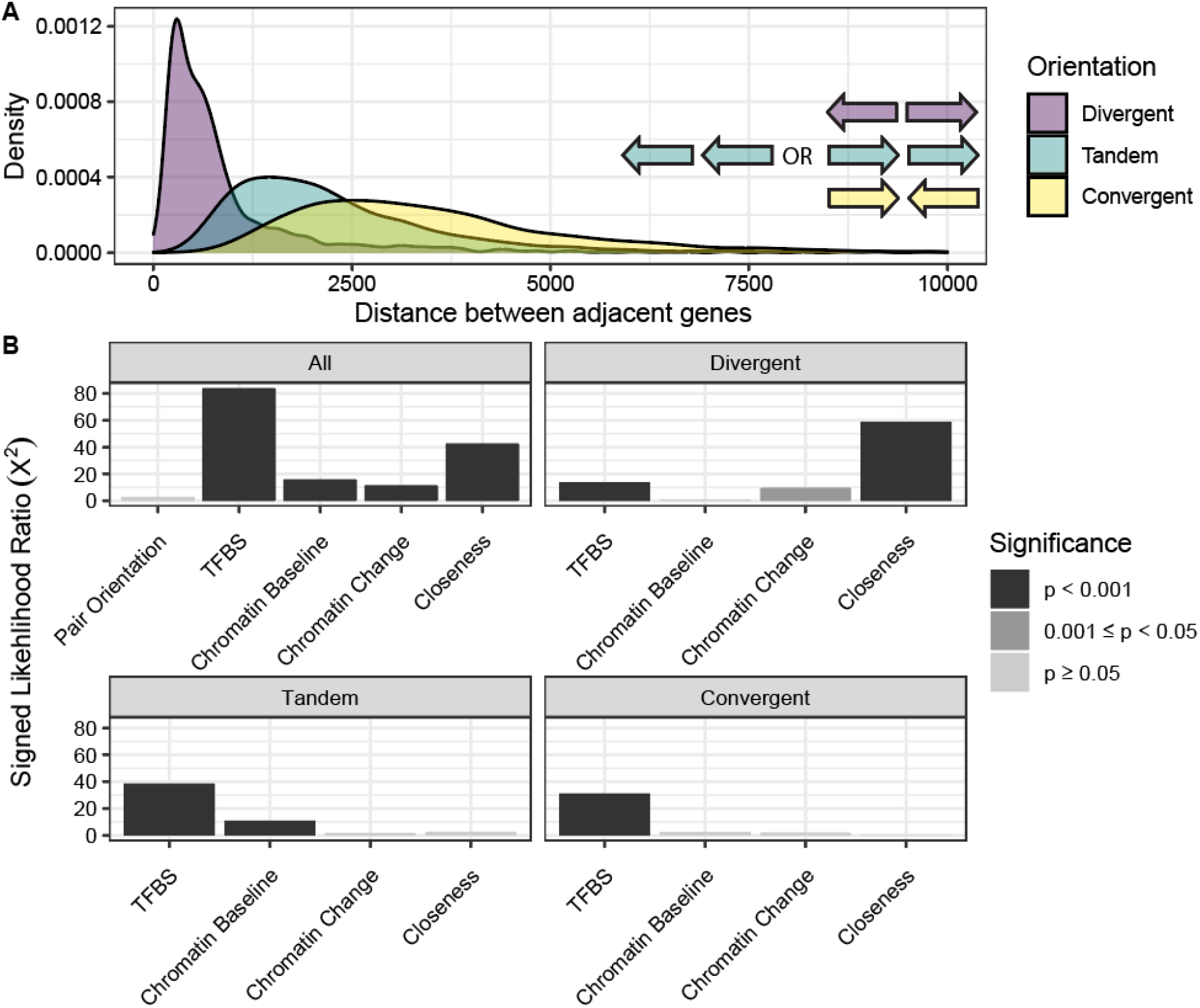
(A) Distribution of intergenic distances for gene pairs in each of the three possible orientations (divergent, tandem, convergent). (B) Influence of various features on paired hotspot effects. Effect directions were plotted based on the sign of the feature’s slope in the linear model; all effects had a positive sign. Pair orientation is plotted with a positive effect.

The features considered above tend to be correlated with each other. For example, divergently expressed genes are close to each other and also share an intergenic region where the same TFBSs may influence their expression. To quantify the contributions of these features relative to each other, we included them as predictor variables in a generalized linear model of the number of paired hotspot effects for adjacent gene pairs. To gauge the significance of a given feature, we compared the full model, which contained all features, to models that did not contain the given feature. Overall, the full model explained a modest (6.4%) but highly significant (p < 2.2×10^−16^) fraction of the variation in paired hotspot effects among gene pairs. Several features had significant effects (Fig 4), but no feature was highly predictive of whether a given gene pair is often affected by hotspots. These modest feature effects are consistent with the magnitude of the correlations reported above, and are expected given the small effects of individual *trans*-eQTLs, which create inescapable noise in our quantification of paired hotspot effects. Nevertheless, these analyses can quantify the importance of features relative to each other, while controlling for correlations among features.

TFBS similarity explained the greatest amount of variation, such that gene pairs with shared TFBSs had more paired hotspot effects (Fig 4B, Suppl Fig 3B). Chromatin baseline similarity and similarity in chromatin change each had smaller but significant effects (Fig 4B). There was no independent effect of gene pair orientation, presumably because the differences between orientations we noted above mostly reflect physical distance (Fig 4A) (Cohen et al., 2000). Across all adjacent gene pairs, physical proximity was the second most important feature, such that genes whose start sites are located closer to each other had more paired hotspot effects (Fig 4B, Suppl Table 3). Importantly, while chromatin baseline similarity (Rho = −0.24; p = 2.2×10^−16^) and chromatin change similarity (Rho = −0.04; p = 0.006), but not TFBS similarity (Rho = −0.02; p = 0.12), are negatively correlated with distance, our model controls for these features, such that they cannot account for the independent effect of distance.

Dividing gene pairs by their orientation showed that the overall distance effect was almost exclusively driven by variation among the divergently oriented gene pairs (Fig 4B). Most divergent gene neighbors are separated by less than 1,000 bp (Fig 4A). This is the same range within which coexpression of adjacent genes is most prominent in yeast, an observation that has been attributed to non-specific, promiscuous interactions between upstream activating sequences and promoters. (Cohen et al., 2000; Quintero-Cadena & Sternberg, 2016). Thus, proximity-based promiscuity of *cis*-regulatory elements could explain why *trans*-eQTL hotspots tend to affect both genes in close, adjacent gene pairs.

In sum, there are three separate reasons for the observation that *trans*-eQTL hotspots tend to affect both members of adjacent gene pairs. First, adjacent genes tend to be regulated by similar transcription factors, which results in paired hotspot effects through vertical mechanisms. Second, for genes in close physical proximity, the effect of a *trans*-eQTL on one gene can also change the expression of the other gene via horizontal mechanisms. Third, similarity in chromatin state, both at baseline and during chromatin dynamics, also increases the probability that adjacent genes tend to be affected by the same hotspots.

### Many adjacent genes with paired hotspot effects do not share a common function

Functionally related genes tend to be clustered along the genomes of humans (Al-Shahrour et al., 2010; Andrews et al., 2015; Caron et al., 2001), yeasts (Cohen et al., 2000; Eldabagh et al., 2018; Pál & Hurst, 2003; Poyatos & Hurst, 2007), flies (Spellman & Rubin, 2002), mice (Li et al., 2005), worms (Kamath et al., 2003), and zebrafish (Ng et al., 2009). This clustering has been suggested to facilitate coexpression of genes in pathways (but see (Kustatscher et al., 2017)). We asked whether paired hotspot effects tend to occur at genes with related functions.

To quantify functional similarity among gene pairs, we used two metrics. First, we computed semantic similarity based on the “biological process” annotations in the Gene Ontology database (Ashburner et al., 2000; The Gene Ontology Consortium, 2019). Second, genes with similar biological function tend to have similar synthetic genetic interaction profiles, as previously quantified (Costanzo et al., 2016). While not all genes have high quality Gene Ontology annotations, the genetic interaction profiles cover the vast majority of gene pairs in a systematic manner. In both metrics, adjacent genes showed modestly but significantly higher functional similarity than non-adjacent genes (semantic similarity: median 0.199 vs 0.195, Wilcoxon test p = 0.002; genetic interaction similarity: median 0.02 vs 0.005, p < 2.2×10^−16^).

We asked how these two measures of functional similarity relate to the molecular features of adjacent genes we compiled above. Genes with higher semantic similarity were more likely to be regulated by similar TFs (p = 6×10^−10^; Suppl Table 2; see also (Hershberg et al., 2005)) and, to a weaker extent, have similar chromatin dynamics (p = 1×10^−5^). By contrast, genetic interaction similarity was not influenced by these two features (p > 0.4). Instead, genes with similar genetic interaction patterns were likely to be physically close to each other (p = 0.0003). In line with these results, semantic similarity and genetic interaction similarity were not correlated with each other (Rho = 0.009, p = 0.6). Thus, these two measures capture different aspects of functional similarity.

We tested the relationship between functional similarity and paired hotspot effects. Genetic interaction similarity was weakly but significantly correlated with paired hotspot effects (Rho = 0.05, p = 0.003), while semantic similarity was not (Rho = 0.02, p = 0.10). Because functional similarity and paired hotspot effects are both shaped by similar sets of molecular features, their correlation could be confounded by these shared influences. To control for this, we computed residuals of each metric, removing the influence of the molecular features. Correlations between the residuals of paired hotspot effects and functional similarity remained weak, and became less significant (genetic interaction similarity: Rho = 0.04, p = 0.02; semantic similarity: Rho = 0.02, p = 0.23). The weak magnitude of these relationships suggests that many adjacent genes with paired hotspot effects do not share a common function.

An example of paired hotspot effects and their relation to gene function is provided by a hotspot that is caused by a non-synonymous variant in the Oaf1 transcription factor (Lutz et al., 2019), which regulates the transcription of genes involved in fatty acid metabolism (Baumgartner et al., 1999). The *OAF1* hotspot affects 39 doublets, a number that was never observed in our permutations, which showed a median (and mean) of 16 doublets and a maximum of 36 doublets. This hotspot alters the expression of genes involved in fatty acid metabolism (Lutz et al., 2019), including many annotated transcriptional target genes of Oaf1 (Bergenholm et al., 2018). For example, the hotspot alters the expression of the beta unit of fatty acid synthetase, *FAS1*, which is transcriptionally regulated by Oaf1. However, the promoters of 74% of genes affected by the *OAF1* hotspot are not known to be bound by Oaf1 (Bergenholm et al., 2018), and the hotspot appears to alter the expression of some of these genes via horizontal mechanisms. For example, the hotspot affects the expression of the *PRS1* gene immediately downstream of *FAS1*. *PRS1* is not a known transcriptional target of Oaf1, and encodes an enzyme involved in nucleotide, histidine, and tryptophan biosynthesis. These processes are not obviously involved in fatty acid metabolism. In summary, horizontal propagation of the effects of *trans*-eQTLs holds the potential of creating unexpected links between the genes that cause a *trans*-eQTL and the genes whose expression they affect.

## Discussion

In this work, we revealed a previously unrecognized mode by which *trans*-acting genetic variation can affect the expression of multiple genes. Typically, *trans*-eQTL effects are thought to propagate through regulatory pathways that are functionally related to the causal eQTL gene. In the alternative mode we describe here, the effects of a *trans*-eQTL can spread along a given chromosomal region and affect potentially unrelated genes (Fig 5).

**Figure 5:**
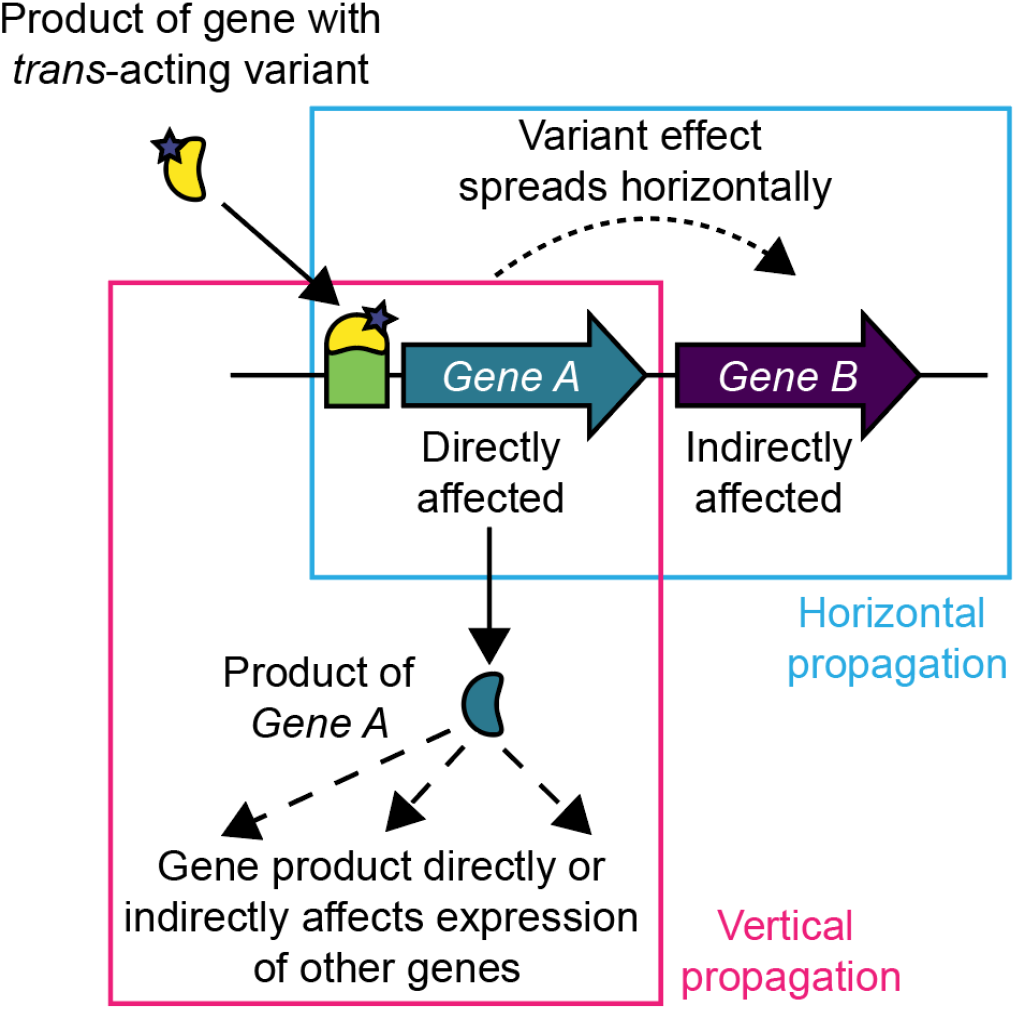
A schematic illustrating two modes by which *trans*-acting sequence variation affects direct and indirect target genes. Left (pink box), the effects of a *trans*-acting variant percolate “vertically” among genes by directly targeting some genes for which altered abundance of the gene product influences other genes in turn. Right (blue box), effects of *trans*-acting variation propagate “horizontally” to genes located proximally to a direct target gene.

We showed that *trans*-eQTL hotspots in a yeast cross affected pairs of neighboring genes more frequently than expected by chance. The effect of the hotspots on adjacent gene pairs tended to involve genes that showed stronger coexpression in response to a broad range of perturbations. Thus, a straightforward explanation for the effects of eQTL hotspots on adjacent genes is that the genetic variants underlying the hotspots trigger the same genomic mechanisms that also shape gene-gene coexpression.

We examined molecular features that could drive pairing of hotspot effects. The strongest determinant was regulation of neighboring genes by similar sets of TFs. This result is consistent with prior observations that genes with similar function, which tend to be regulated by similar TFs as part of transcriptional regulons, tend to be physically colocated in the yeast genome (Hershberg et al., 2005). Pairing of hotspot effects due to regulation by similar TFs is expected under conventional, vertical propagation of hotspot effects in a pathway, if pathway members are located next to each other in the genome. However, the second strongest determinant of paired hotspot effects was the proximity of adjacent genes on the chromosome. Crucially, this analysis controlled for shared, vertical regulation by similar TFs. Thus, *cis*-acting mechanisms that extend horizontally along a chromosome in a manner that is dependent on physical distance must be responsible for this effect.

A possible mechanism involves chromatin states that extend across adjacent genes, as suggested by studies of individual gene pairs (Batada et al., 2007; Ebisuya et al., 2008; Raj et al., 2006)). Indeed, adjacent genes that had correlated chromatin marks in normally growing cultures and during response to cellular stress were more likely to be affected by hotspots as a doublet. This relationship suggests that shared chromatin state is one mechanism by which hotspot effects spread horizontally.

Proximity-dependent promiscuous effects of *cis*-regulatory elements are another possible mechanism that could result in paired hotspot effects. In eukaryotes, transcriptional regulation is shaped by *cis*-regulatory elements bound by sequence-specific transcription factors (Wittkopp & Kalay, 2012). These elements, which include yeast upstream activating elements and enhancers in higher eukaryotes, interact with the core promoters of specific genes (Hahn & Young, 2011). Under proximity-dependent promiscuity, which has also been called “enhancer-promoter (EP) theory” (Cera et al., 2019; Quintero-Cadena & Sternberg, 2016), *cis*-regulatory elements can also regulate transcription from any promoter within a certain physical distance (Butler, 2001; Ebisuya et al., 2008). This non-specific activity decays as distance between the element and a gene increases. Distance-dependent coexpression has been described in species ranging from yeast to human, and the reach of this phenomenon scales as a function of genome size (Quintero-Cadena & Sternberg, 2016). In the compact yeast genome, distance-dependent coexpression is detectable within a range of about 1,000 bp. In our data, distance only influenced pairing of hotspot effects among divergently expressed gene pairs. Most of these pairs are located less than 1,000 bp apart, well within the reach of non-specific EP processes. Thus, pairing of hotspot effects may be partly driven by proximity-dependent promiscuity of *cis*-regulatory elements.

While the precise mechanistic basis of horizontal propagation of hotspot effects remains unclear, its potential implications are profound. Under this model, a *trans*-eQTL first alters the expression of a distant gene via conventional, vertical regulation. Altered expression of the target gene causes molecular changes that ripple outward along the chromosome and change the expression of other genes in the same area. This horizontal extension of hotspot effects has the potential to create unexpected links between causal DNA variants and complex traits. If the *trans* effect extends to a neighboring gene with a function that is unrelated to the causal gene in the hotspot, the altered expression of this neighboring gene could alter cell physiology in ways that are not obvious from the function of the causal eQTL gene. Such indirect consequences could contribute further complexity to the genetic basis of quantitative traits (Boyle et al., 2017).

Compared to yeast, our knowledge of human *trans*-eQTLs remains limited (Aguet et al., 2017; Battle et al., 2014; Brynedal et al., 2017; Fehrmann et al., 2011; Grundberg et al., 2012; Heinig et al., 2010; Lee et al., 2014; Small et al., 2011; Wright et al., 2014; Yao et al., 2017). Nevertheless, the prevalence of EP effects across species including humans (Quintero-Cadena & Sternberg, 2016) means that horizontal spreading of *trans*-effects is expected to also exist in other species.

## Materials and methods

Code and datasets used in this work are available at (https://github.com/Krivand/Trans-acting-genetic-variation-affects-the-expression-of-adjacent-genes). All figures, intermediary processing objects, tables, and analyses can be generated directly from the code on GitHub by following the instructions at the top of the R script. Figures in this table were paneled and combined in a separate image processing package.

Unless otherwise specified, analyses were conducted in R 3.6.0 (R Development Core Team, 2010), using the following packages. C++ code was run using Rcpp 1.0.4 (Eddelbuettel & François, 2011). Data processing was performed using tidyverse 1.3.0 (Wickham et al., 2019). Genomic range data were imported using Rtracklayer 1.46.0 (M. Lawrence et al., 2009) and processed using Genomic Ranges 1.38.0 (Michael Lawrence et al., 2013) and Biostrings 2.54.0 (H. Pagès, 2017). Plot color schemes were provided by the Viridis color package originally developed as part of Matplotlib (J. D. Hunter, 2007).

### Datasets

All datasets used in this work are detailed in Suppl Table 3.

### Gene annotations

Gene models were obtained from yeastmine (Balakrishnan et al., 2012; Fisk et al., 2006). The BY and RM strains differ in the presence or absence of stretches of genes close to the ends of the chromosome ends. *trans*-eQTL effects on these subtelomeric genes may appear to be highly correlated simply due to their shared presence or absence in the BY / RM segregants. Therefore, we removed subtelomeric genes, defined as all genes whose start or stop codon was located outside of the genetic markers forming the linkage map in Source Data 3 from (Albert et al., 2018). We retained only “verified open reading frames” (ORFs) (Fisk et al., 2006) because some genes not annotated to be in this category (such as “dubious” ORFs or noncoding genes) overlap verified ORFs (sometimes on the antisense strand), complicating analyses of intergenic distances and TFBS annotations.

### Hotspot effects

The effects of hotspots were represented in the “hotspot matrix”. This matrix was generated from previously published data (Source Data 9 from (Albert et al., 2018)), which cataloged the effects of all 102 hotspots (columns) on 5,629 expressed genes (rows). We removed 734 genes which were not verified ORFs (Fisk et al., 2006) and introduced a row of zeros for 17 genes that were expressed in the segregant data but that were not affected by any of the 102 hotspots. Genes were defined as expressed if they were present in the log_2_(TPM) expression value table in Source Data 1 from (Albert et al., 2018). The rows of the hotspot matrix were ordered by chromosome and the position of the genes on each chromosome. We removed subtelomeric genes and retained only verified ORFs as described above. After these filters, the hotspot matrix contained 4,912 genes, of which 4,878 were affected by at least one hotspot.

### Doublet definition

A doublet was defined as two adjacent genes with non-zero effects of equal sign from a given hotspot. A single gene can be involved in up to two doublets, one with each of its adjacent genes. Genes at the ends of chromosomes can only be involved in one doublet. For each hotspot, we counted the number of doublets among the genes affected by the hotspot.

### Permutation Tests

To assess whether the observed doublet count exceeded random chance, we used a permutation test. For each hotspot, we performed 10^5^ permutations, in which the given hotspot effects were randomly shuffled across the genome. In each permutation, we counted the number of doublets.

To determine p-values, we counted the number of permutations with a number of doublets greater than or equal to those observed in the real data for the corresponding hotspot. Multiple testing correction across 102 hotspot was performed by dividing the significance threshold of p = 0.05 by 102. This process was repeated for doublets excluding divergent genes, as well as for triplets, quadruplets, quintuplets, and sextuplets.

### Coexpression matrices

Expression datasets (Suppl Table 3) were ordered by chromosome and gene order, with genes in rows and perturbations in columns. We universe-normalized each dataset by removing genes not present in the BY / RM expression dataset (Source Data 1 from (Albert et al., 2018)) and by adding a null-row for any genes that were present in the BY / RM expression dataset but absent from the given expression dataset. Quantile normalization was performed using the normalize.quantiles function from preprocessCore (Bolstad, 2017), and used to generate Spearman correlation matrices via Hmisc’s Rcorr function (Harrell, 2020). The datasets from Albert & Bloom *et al*., 2018, Brem & Kruglyak 2005, Fleming *et al*. 2002, Hughes *et al*. 2000, and Schurch *et al*., 2016 were treated similarly, with the following changes:

Fleming *et al*. 2000 and Hughes *et al*. 2000: Data had been preprocessed following the procedures described in (Huttenhower et al., 2006). Universe normalization and quantile normalization were performed as above.

Brem & Kruglyak 2005: Data had been preprocessed following the procedures described in (Huttenhower et al., 2006). We used the mean of both microarray dye swaps as the expression measurement for each gene. Universe normalization and quantile normalization were performed as above.

Schurch *et al*. 2016: Processed counts were obtained from GitHub (https://github.com/bartongroup/profDGE48/blob/master/Preprocessed_data/WT_countdata.tar.gz). Raw data from which these processed counts were derived is available at the European Nucleotide Archive repository (http://www.ebi.ac.uk/ena/data/view/ERX425102) under the accession number PRJEB5348, where they had been deposited by (Gierliński et al., 2015). Counts for each gene were universe-normalized as above and filtered to retain only genes with at least a single read in at least half of the samples. Read counts were normalized by dividing them by the length of the gene. A half-count was added to each count before taking the natural log of each value. Counts were centered within each sample by subtracting the mean expression of all expressed genes in the sample from all expression values in that sample. Data then underwent quantile normalization as above.

Albert & Bloom *et al*., 2018: Log(TPM) reads in Source Data 1 from (Albert et al., 2018) were processed to remove the effect of genetic variation, processing batch, and growth phase. As a proxy for the growth phase, we used the optical density at the time the cultures had been collected (Source Data 2 from (Albert et al., 2018)). For each gene, we built a fixed-effects linear model using processing batch, optical density, and peak markers of each significant *cis*- and *trans*-eQTL reported for this gene (Source Data 3 from (Albert et al., 2018)). The residuals from these models were centered by subtracting the mean of all expression measurements for that segregant from all expression values for that segregant. We then performed quantile normalization as above. To gauge if removal of eQTL effects had been successful, we examined genes with strong eQTLs. Prior to removal, such eQTLs cause the expression values of their target gene to show a bimodal distribution. We examined the gene *STE2*, which is affected by a strong *trans*-effect arising from the mating type locus, and the gene *HO*, which is affected by a strong local effect caused by an engineered deletion of this gene in the RM strain. Both genes had bimodal distributions prior to correction, which became unimodal in the residuals of their respective linear models (Suppl Fig 4).

### Comparison of similarity matrices

The various similarity matrices were compared using Spearman rank-based correlations between the upper triangles of the two given matrices.

### Quantifying paired hotspot effects across hotspots

To capture the degree to which both genes in a given pair were affected by hotspots, we constructed a matrix containing hotspot effect summaries. We initially set every possible gene pair to a value of zero and then examined the 102 hotspots. If a hotspot affected both genes in a pair in the same direction, the score was increased by one. If both genes in a pair were affected in different directions, the score was decreased by one. Thus, these summarized hotspot effects are able to capture gene pairs that are in repulsion, such that they tend to be affected by multiple hotspots in opposite directions. We chose this metric for comparison to coexpression because it was able to capture negative values for gene pairs with anticorrelated expression, which may also show pairing of hotspot effects with opposite direction.

In our linear models of possible mechanisms underlying paired hotspot effects, we used a matrix that counted the number of hotspots affecting both members of a pair in the same direction. While this metric ignores pairs of genes with hotspot effects in opposite direction, we chose it for two reasons: 1) the absence of negative values allowed us to use a count-based generalized linear model (see below), and 2) many of the features we used to model paired hotspot effects cannot take on or are not meaningful for negative values (e.g. TFBS similarity, or distance between genes).

### Shared regulation of genes by transcription factors

We used a curated collection of transcription factors annotated to regulate each gene in the yeast genome (Monteiro et al., 2020). These data were binary, with a value of zero indicating that a given TF is not known to regulate a given gene, and a value of one indicating that it does. TF profile similarity between gene pairs was calculated using the Jaccard index (Veerla & Höglund, 2006).

### Nucleosome occupancy measures

We used two nucleosome occupancy datasets: an ATAC-seq dataset (Schep et al., 2015) and a dataset based on a chemical cleavage assay (Chereji et al., 2018). Specifically, we used the nucleosome occupancy files available at the NCBI Gene Expression Omnibus (GEO; http://www.ncbi.nlm.nih.gov/geo/) under accession numbers GSE66386 and GSE97290, respectively. Data had been pre-processed with scores indicating low to high nucleosome occupancy for a given genomic range. For the chemical cleavage data, means for occupancy at each base pair across three replicates was used for our calculations.

In each dataset, we calculated two metrics. 1) Mean occupancy across adjacent gene bodies was defined as the mean nucleosome occupancy of the region from 500 bp upstream of the first gene to 500 bp downstream of the second gene. 2) Similarity in occupancy of adjacent genes was computed as the log of the inverse of the difference in mean nucleosome occupancy score of both gene bodies.

### Similarity in chromatin marks at baseline and during stress response

Chromatin similarity at baseline and change in chromatin similarity were calculated using ChIP-seq data published by (Weiner et al., 2015). Specifically, we used the nucleosome maps (Table S2) and normalized marks (Table S3) available separately at (https://www.cell.com/molecular-cell/fulltext/S1097-2765(15)00094-5) or combined at the NCBI Gene Expression Omnibus (GEO; http://www.ncbi.nlm.nih.gov/geo/) under accession number GSE61888. Table S2 from Weiner et al. contains the mapped positions of 66,360 nucleosomes in wild-type yeast (strain BY4741) and Table S3 from Weiner et al. contains the levels of all 26 chromatin modifications normalized to the data in their Table S2. Thus, the values in their Table S3 represent estimates of the log ratio of ChIP coverage with respect to each sample input, with quantile normalization performed within the time series for each chromatin mark.

We calculated similarity in baseline chromatin state between a pair of genes as the spearman correlation between the normalized levels of the 26 chromatin marks on the +1 nucleosomes of the two genes in mid-log growth, before the application of the stressor. Similarity in chromatin change was calculated as the spearman correlation between the difference in normalized levels of the 26 chromatin marks on the +1 nucleosomes of neighboring genes between baseline and 15 minutes after the cells were exposed to diamide.

### Features influencing the number of hotspots that affect gene pairs

To gauge the relative importance of various features on paired hotspot effects, we fitted a negative binomial generalized linear model, using the glm.nb() function from MASS 7.3-51.6 (Venables et al., 2002). The response variable was the number of times adjacent gene pairs were affected by the same hotspot in the same direction. Pairs with zero paired effects were omitted. Some genes were affected by more hotspots than others, which trivially increases the number of times they could form a doublet with their adjacent gene. To control for this, we included two exposure variables as offsets, each giving the natural log of the number of times a given gene in a pair was affected by hotspots.

As predictor variables, our models included: 1) the orientation of genes with respect to each other (i.e. convergent, tandem, or divergent), 2) the similarity in how gene pairs were regulated by transcription factors, 3) physical “closeness” on the chromosome, which was the natural log of the inverse of distance in bp, 4) similarity in chromatin marks between +1 nucleosomes, and 5) similarity in cell stress-induced changes in chromatin marks between +1 nucleosomes. To calculate the significance of each of these features, we fit a full model with all predictors, as well as reduced models that omitted the given feature. Model comparison between the full model and each reduced model was performed using type III anova as implemented by the Anova() function in the car 3.0-8 package (Fox & Weisberg, 2019). We also fit equivalent models to subsets of the data, separately for each of the three possible gene orientations.

### Functional similarity between genes

#### Gene ontology method

We used the GoSemSim package (Yu et al., 2010) with the Wang method (Wang et al., 2007) and best match average scoring to deal with the semantic similarity scores of multiple GO biological process terms. GO data were obtained via the org.Sc.sgd.db 3.10.0 package (Carlson, 2017).

#### Genetic interaction similarity method

These data were preprocessed and derived from Pearson correlations provided at (https://thecellmap.org/costanzo2016/ (Costanzo et al., 2016)). This matrix was universe-normalized by removing any columns and rows corresponding to genes not present in the BY / RM data (Albert et al., 2018) and adding null columns for genes present in the BY / RM data that were not present in the interaction dataset.

### Genes regulated by Oaf1

All genes directly regulated by Oaf1 were obtained from previously published data ((Bergenholm et al., 2018) DATA SET S1). Genes were considered to be regulated by Oaf1 if their promoters were bound by Oaf1 in any of the four conditions studied by (Bergenholm et al., 2018).

## Acknowledgements

We thank Sheila Lutz and Christian Brion for critical feedback, and Kristine Trotta as well as members of the Albert lab (Randi Avery, Mahlon Collins, Kelsey Johnson, and Kaushik Renganaath) for comments on the manuscript.

## Supplementary Material

Supplementary Table 1: Summary statistics for hotspot permutations, real doublet counts, and p-values. P-values listed as “0” indicate that no count greater than or equal to that of the real data was observed in the permutations. Thus, while the corresponding p-value is <1×10^−5^, we report these cases as “0” to avoid Excel formatting cells with a “<“ sign as text.

Supplementary Table 2: Output of type III ANOVAs for generalized linear models.

Supplementary Table 3: Datasets used in this work, with details on the source of data presented here.

## Supplementary Figures

**Supplement Figure 1:**
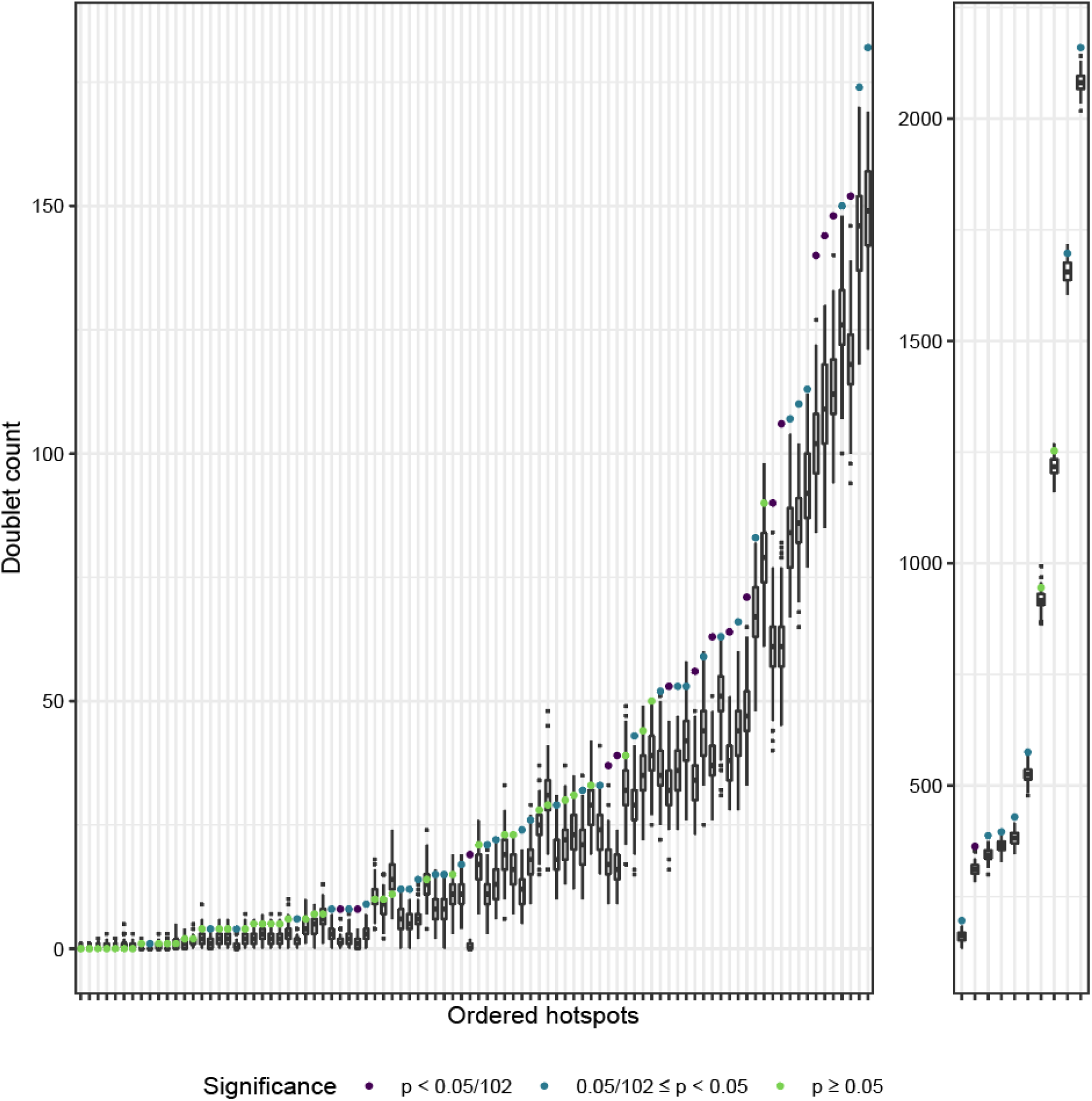
The number of doublets in real data (colored circles) are plotted against the number of doublets in permuted data (white boxes). Boxes extend from the 25th to 75th percentile. Whiskers extend from each box to the largest or smallest values within 1.5 times the difference in the 25th to 75th percentile. Data beyond the whiskers are considered outliers and plotted as small black squares. Circles are colored by the number of permutations in which a number of doublets was counted that was greater than or equal to the number observed in the real data. Hotspots are ordered along the x-axis based on the value of the doublet count in the real data. For clarity, the ten hotspots with the largest number of doublets are shown with a different scale.

**Supplementary Figure 2:**
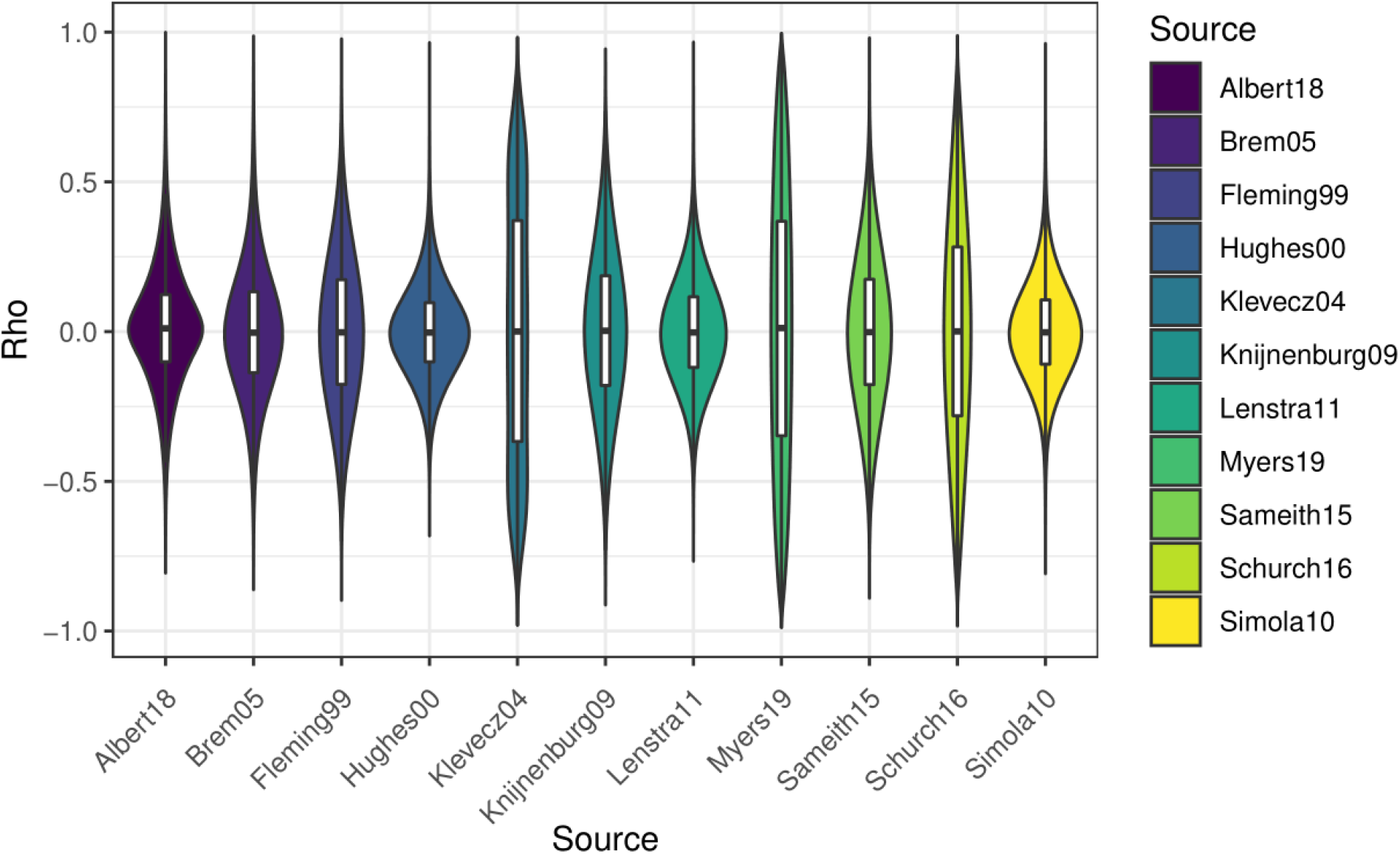
The distribution of Rho values for each coexpression matrix generated from the 11 expression datasets.

**Supplementary Figure 3:**
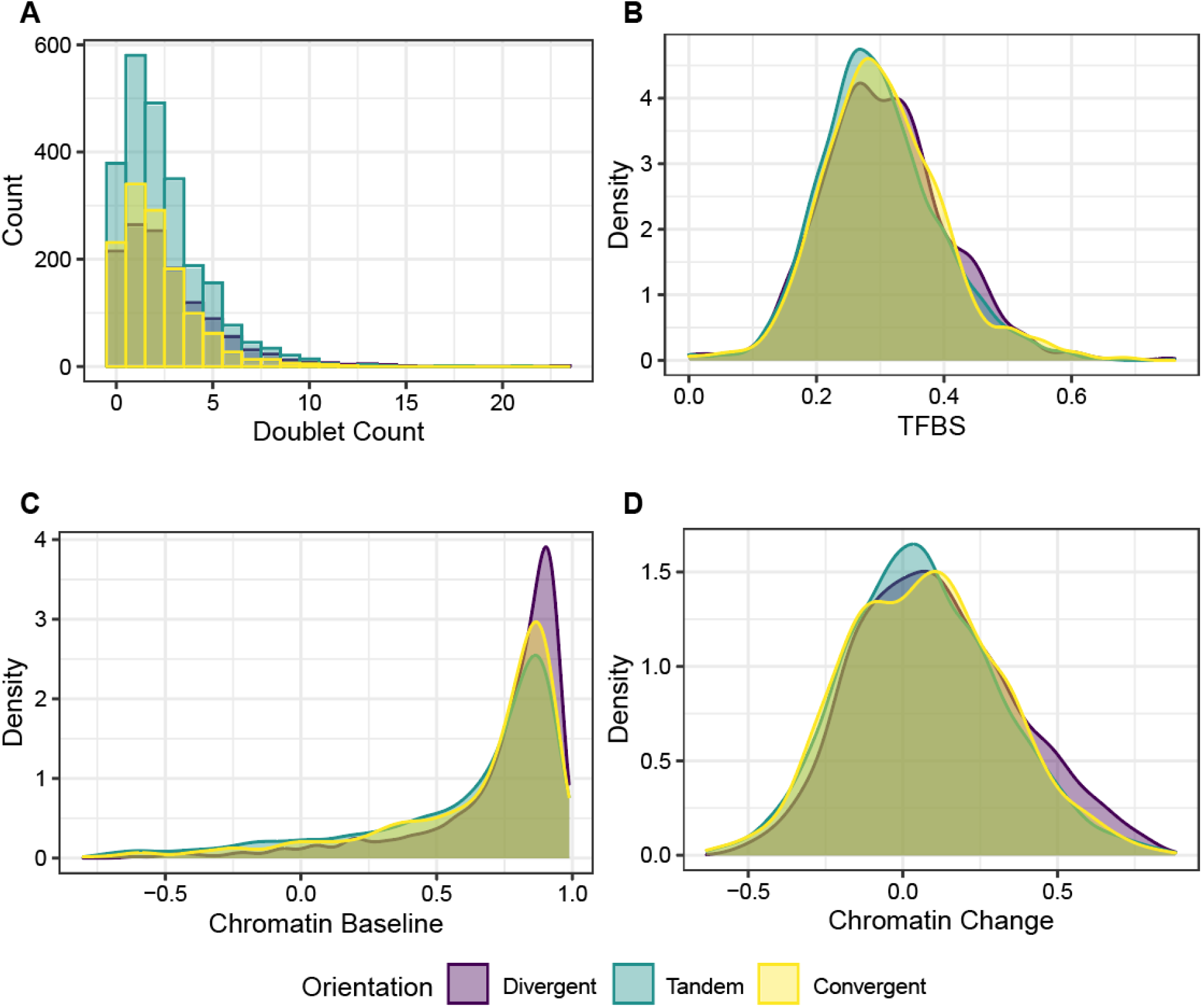
Distributions of doublet counts per neighboring gene pair and predictor variables, separated by gene pair orientation. Distance is shown in Fig 4A. (A) Number of times gene pairs were targeted by the same hotspot. (B) Similarity in regulation by transcription factors. (C) Correlation between marks on +1 nucleosomes without perturbation. (D) Correlation in change in marks in response to cell stress on +1 nucleosomes.

**Supplementary Figure 4:**
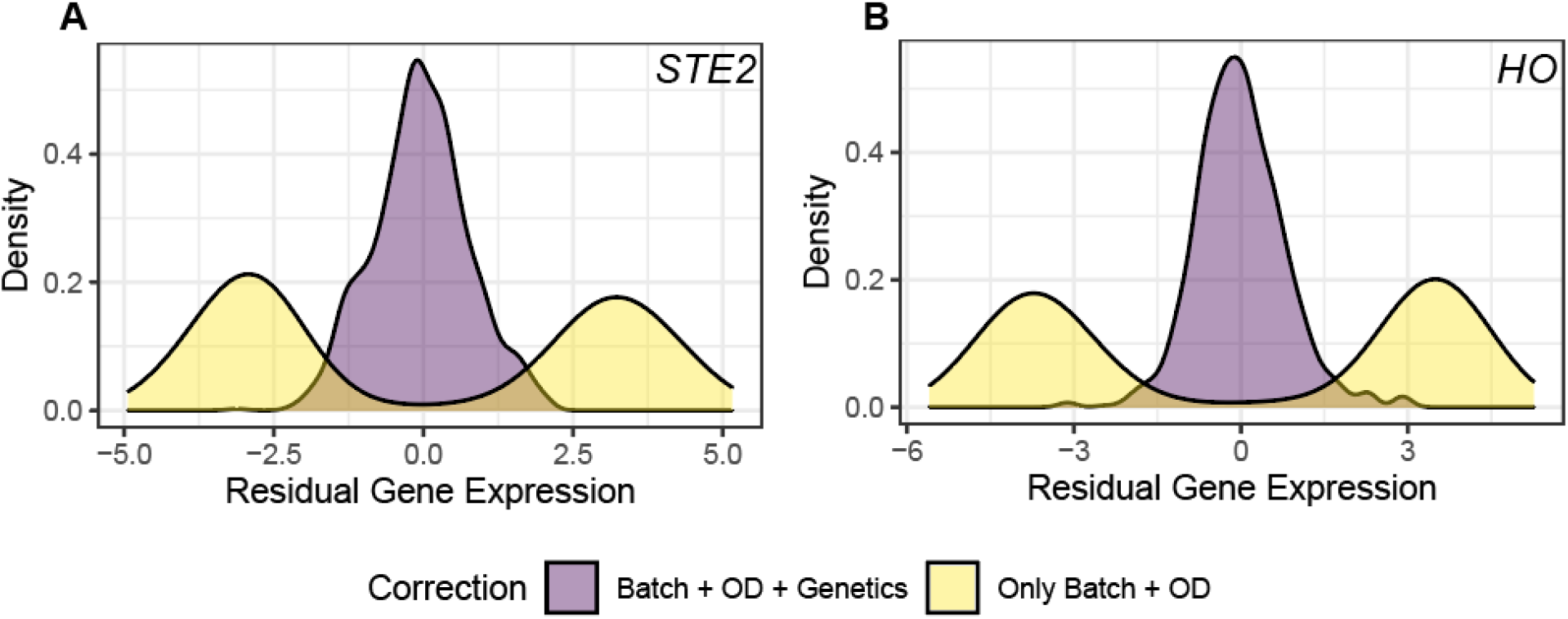
Distribution of residual gene expression for two genes after correction for either only batch and optical density (OD) or batch, OD, and genetic variation (A) Residuals for *STE2*, whose expression is strongly affected by the mating type locus before correction for genetic variation. (B) Residuals for *HO*, which is deleted in the RM but not the BY strain used in Albert & Bloom *et al*., 2018, resulting in a strong local eQTL in the BY / RM progeny.

